# Global, single-cell-resolution of antiviral defenses in marine prokaryoplankton

**DOI:** 10.64898/2026.09.15.751803

**Authors:** Alaina R. Weinheimer, Julia Brown, Greg Gavelis, Tianyi Chang, Miglė Štitilytė, Giedrius Gasiunas, Ramunas Stepanauskas

**Affiliations:** Bigelow Laboratory for Ocean Sciences, East Boothbay, USA; Helmholtz Institute for Functional Marine Biodiversity, 26129 Oldenburg, Germany; Gensinta, Vilnius, Lithuania; CasZyme, Vilnius, Lithuania; Institute of Biotechnology, Life Sciences Center, Vilnius University, Vilnius, Lithuania; University of Aalborg, Aalborg, Denmark

**Keywords:** antiviral defenses, microbial ecology, marine microbiology, single-cell genomics, prokaryoplankton, ocean microbiome, deep ocean

## Abstract

**Background:** Growing evidence suggests that variation in resistance to viral infections by marine prokaryoplankton enables the coexistence of both viral and host populations. Recent experimental work on model organisms has revealed dozens of novel antiphage defense mechanisms. This diversity has prompted the hypothesis of the pan-immunity model, in which diverse defenses are a shared resource among closely related individuals to broaden the population’s resistance and minimize an individual’s burden. The composition and abundance of such defense pools in natural microbial communities, however, remain largely unknown. Here, we begin to parameterize resistance in marine prokaryotes and define their defense repertoire by quantifying antiviral defenses across thousands of randomized single-amplified genomes (SAGs) from a global collection of seawater samples.

**Results:** Prokaryoplankton SAGs contained an average of 1.1 defenses, with dark ocean (200 m – 10 km) prokaryotes having a slightly higher genomic load of defenses as compared to the sunlit, surface ocean. The number of defenses per cell was taxon-specific, with little or no relationship to depth, taxon abundance, estimated maximal growth rate, or viral infection rate. The most numerous and taxonomically widespread defenses were restriction modification systems, followed by dGTPase (mostly in Pelagibacterales) in the sunlit ocean and AbiU (mostly in Nitrososphaerales) in the dark ocean. Most other defense types were rare (< 0.1% SAGs) yet distributed across distant taxa (> 3 phyla). We found evidence for the cross-domain exchange of the most common defense system, RM II, between archaea and bacteria, which improves our understanding of the biology of this prevalent defense and could be consequential in predicting target motifs for connecting viruses with potential hosts in epigenetic studies.

**Conclusions:** Collectively, these results suggest lineages restrict genomic real estate for defenses yet enable extensive lateral transfer, potentially for the maintenance of defense variation at the community level rather than only among closely related individuals as suggested by the pan-immunity model. This study provides a quantitative atlas of prokaryoplankton immunity toward grounding our understanding of virus-microbe interactions in the Earth’s largest biome, the open ocean.

## Background

In the open ocean, viruses outnumber microbes by roughly ten-fold^1^. They can short-circuit the movement of nutrients up the marine food web by infecting and lysing microbial cells, a process called the viral shunt ^1^. Roughly 4-20% of prokaryotic cells are estimated to be lysed by viruses in the ocean per day ^2,3^. Growing evidence from culturing, modelling, and metagenomic studies suggests that variation in resistance to infections by prokaryotes contributes to the composition and persistence of both prokaryotic and viral populations ^4–6^. Within the past decade, research on antiviral defenses in model prokaryotes like *Escherichia coli* has vastly increased the number and diversity of known defense systems ^7,8^, with over 150 described at the time of this study, and roughly five defenses found per genome in RefSeq ^9^. The “pan-immunity” model ^10^ has been proposed to explain the maintenance of this defense diversity. According to this model, defense genes with varying specificities and fitness trade-offs are encoded by closely related individuals (cells) to reduce an individual’s defense cost and, collectively, broaden the population’s protection. Whether and how this model impacts virus-host interactions in natural, mixed microbial communities, however, remains unknown.

Experimental work and metagenomic analyses have shown that different defense strategies are enriched under different ecological conditions ^11–13^. Previous studies have found that marine-associated prokaryotes are particularly depleted in defenses relative to other environments (e.g. animal-associated, soil, hot springs, etc.) ^12,14^. However, the variable presence of defense system genes across closely related cells, in addition to the microdiversity of marine prokaryotic genomes overall ^15,16^, may prevent defense genes from assembling in bulk-sequencing approaches altogether ^17^. Additionally, these bulk sequencing approaches do not enable direct linking of a cell with a defense, limiting quantitative inferences. Single cell genomics offers an alternative to bulk sequencing ^16,18^. Cells are physically separated from one another, and their genomes are amplified, sequenced, and assembled into single-amplified genomes (SAGs), thereby providing cell-level resolution of genomic diversity. Furthermore, random sampling of cells can enable a quantitative representation of the microbiome ^19^.

Here, we identified and quantified defenses in thousands of SAGs from prokaryoplankton cells randomly sampled from the sunlit open ocean (< 200 m) within tropical and subtropical latitudes of the Global Ocean Reference Genome (GORG) Tropics collection ^20^. We also contrasted these findings against thousands of SAGs from the global deep ocean (200 m - 11,000 m) of GORG Dark ^21^. The similar methods and scale of GORG Tropics and GORG Dark enabled the exploration of the distribution of defense strategies across depths. We hypothesized that the biogeochemical disparities between the sunlit and dark ocean (e.g., differences in productivity, ratio of viruses to microbes, and taxonomic diversity) would result in distinct defensive arsenals in the prokaryotes that live in these environments ^2,11,14,22^.

## Results and Discussion

### Abundance of defenses across marine prokaryoplankton

We determined the overall distribution of defenses across the sunlit ocean in the GORG Tropics^20^ and dark ocean in the GORG Dark^21^ SAG collections, of which we focused on a subset of SAGs from both collections that were sorted with a Syto9 DNA stain for prokaryotes and contained at least 50% genome completeness (Supplemental Table S1, S2). This resulted in 3,880 SAGs of the GORG Tropics and 2,156 SAGs of the GORG Dark collections. We detected genes of complete defense systems with the tools DefenseFinder^9^ and PADLOC^23^ (Supplemental Table S1, S2, S3). We identified a complete defense system in 69% of SAGs, suggesting that roughly every cell contains a defense when accounting for average genome completeness (67.5%) (Supplemental Fig. S1). The number of defenses per SAG ranged from 0 to 18 with an average of 1.1. To compare to defense recovery in metagenomics, we examined a complementary set of marine MAGs (minimum completeness 50%; see Methods, Supplemental Table S4, S5), using the same detection workflow (Supplemental Table S5). We detected a slightly higher average number of defenses per MAG, 1.4; however, this higher average was driven by a subset of MAGs that contained many defenses (5+) (10% of MAGs versus 0.5% of SAGs) because the frequency of detecting a defense per MAG (45.1%) was lower than that of SAGs (69.3%) (Fig. 1ab; Supplemental Fig. S1). The lower detection rate of defenses in MAGs may be a result of defense genes failing to assemble, as well as the capture of different lineages than SAGs, resulting in more MAGs with 5+ defenses ^16^. Compared to reference prokaryotic genomes, the average number of defenses per GORG SAG is also lower than those in RefSeq, where prokaryotes encode and average of 5 defenses per genome (Fig. 1c) ^9^. This discrepancy between marine prokaryotes and RefSeq could be due to both methodological and biological factors. In terms of method, marine microbes are underrepresented in cultures and experimental work, and are therefore likely to have uncharacterized defenses. In terms of biology, marine microbes live in an oligotrophic habitat, and are thus subject to extensive genome streamlining ^8^.

**Fig. 1.**
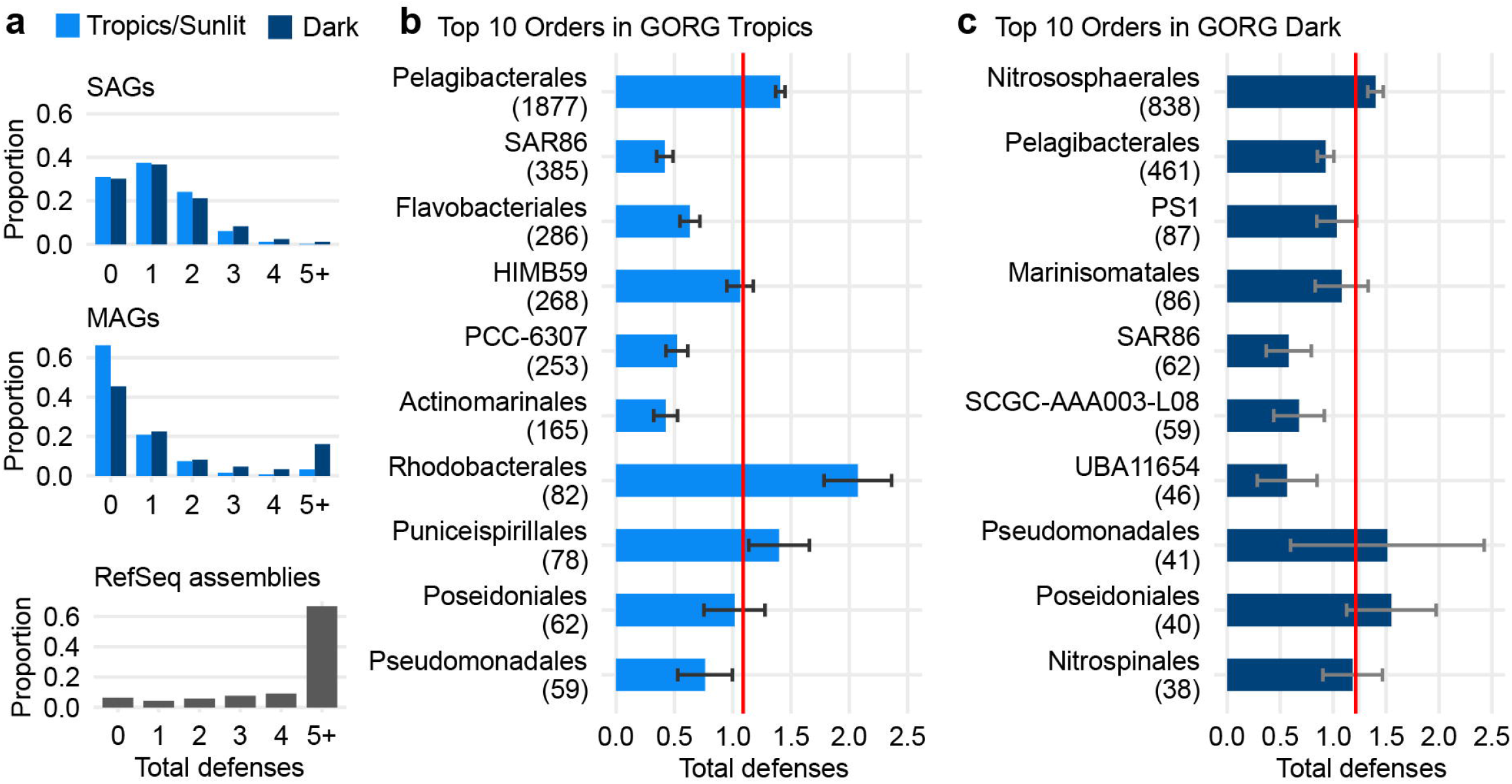
Defense loads across marine and reference prokaryotes. (**a**) Proportion of SAGs, MAGs, or RefSeq assemblies that have a given number of defenses. Colored bars indicate ocean depth within a dataset. in GORG Tropics (light blue) versus Dark (navy) (**b,c**) Average total defenses found per Order in the top ten orders of the GORG Tropics (b) and Dark (c) with number of SAGs in parentheses. Red line indicates dataset mean. Error bars correspond to two standard errors from Order mean.

Between ocean depths, GORG Dark SAGs collectively contained significantly higher defense loads (defined as: number of defense systems and coding sequence density of defense genes) than Tropics SAGs (two-tailed Wilcoxon test *p values* < 0.05; Fig. 1ab; statistics in Supplemental Table S6). Within lineages that contained at least five SAGs in each depth, this depth trend held true for the few significant differences observed at all taxonomic levels, except for lineages that included *Pelagibacter* (Supplemental Table S7, S8). Lineages containing *Pelagibacter* (Proteobacteria, Alphaproteobacteria, Pelagibacterales, Pelagibacteraceae) had higher defense loads in the Tropics than Dark, but there were no significant differences in Proteobacteria when *Pelagibacter* were excluded (Supplemental Tables S7, S8). This general enrichment in defenses in GORG Dark prokaryotes counters predictions that less productive environments with lower viral abundances have fewer defenses ^12,14^. Nonetheless, the variation in defense loads is minimal between the sunlit and dark ocean, with roughly ∼1 defenses per SAG on average in both GORG Tropics (1.10) and Dark (1.21). Perhaps differences in ecological factors between the sunlit and dark oceans may not be strong enough to manifest in substantial differences of defense arsenal size.

### Lineage-specific defense loads

Some lineages had significantly lower defense loads while others were significantly enriched relative to the community mean (Fig. 1de; Supplemental Table S9 for other levels; minimum 5 SAGs to test; two-sided Mann Whitney U test), and we thus examined genomic and ecological factors that might account for these differences. We hypothesized that a lineage’s average genome size or number of genes correlated positively with number of defenses or defense genes, as found in previous surveys ^24,25^, but neither of these relationships were significant across all taxonomic levels (Phylum to Species) (Supplemental Table S10; Order level in Supplemental Fig. S2ab). Ecological features could also select for higher defense investment, since lineages that are (i) more abundant or (ii) slower-growing could be more vulnerable to encountering a virus. We found neither of these factors correlated significantly with a higher defense load (Supplemental Table S10; Supplemental Fig. S2cd). We also estimated viral interaction rates of lineages based on the percent of SAGs that contained viral genomic material (Supplemental Table S1, S11). This estimated infection rate did not significantly correlate with defense loads (Supplemental Table S10; Fig. S2e). However, the variation in number of defenses and variation in defense coding sequence density per lineage decreased as the abundance of a lineage increased, suggesting a convergence on a lineage-specific number of defenses (Supplemental Table S10; Supplemental Fig. S2f).

We then looked individually at two particularly important class-level lineages in the sunlit ocean, alphaproteobacteria and cyanobacteria. In GORG Tropics, alphaproteobacterial lineages tended to have equal or higher defense loads than the community mean, while lineages like Cyanobacteria order PCC-6307 tended to have fewer defenses (Fig 1de). Alphaproteobacteria are known for encoding gene transfer agents and prophages that can be hotspots of defenses ^26,27^; but less than 1% of defenses detected here were also encoded on a contig with a viral hallmark gene to suggest this association (Supplemental Table S11). Meanwhile, marine cyanobacteria are known for encoding fewer defenses than other lineages ^28^, which supports our observations here. Taken together, our results suggest that defense load in prokaryotes largely reflects phylogenetic constraint, and is not easily explained by life history (e.g. abundance, doubling time, genome size) or ecology (e.g. depth, infection rate).

### Diversity of defense systems in marine prokaryoplankton

Of the 146 known system types and 332 subtypes examined, we found that GORG SAGs contained 30.7% (58) of these systems and 29.5% (98) of these subtypes. The subtypes were defined by DefenseFinder^9^ and reconciled with PADLOC^23^ system labels (Supplemental Table S3). Most (61%) of the systems were found in both the GORG Dark and Tropics SAGs, with 36% (21 systems) only found in the GORG Dark and 3% (2 systems) only found in the GORG Tropics. The systems unique to each dataset were too rare (1 - 2 SAGs), to determine whether their abundance significantly differed between water column depths. Restriction modification (RM) systems were the most abundant system type in both GORG Tropics and Dark (Fig. 2), found in 51% and 47% of SAGs, respectively. Furthermore, 85% of GORG phyla contained at least one RM system, highlighting the well-known conservation of this system in prokaryotes ^9,14^. The most common RM subtype in both GORG Tropics and Dark was subtype II (Fig. 2), present in 42% and 29% of SAGs, respectively. Subtypes I, IIG, III, and IV were also common but differed in prevalence between GORG-Tropics and GPRG-Dark (Fig 2).

**Fig. 2.**
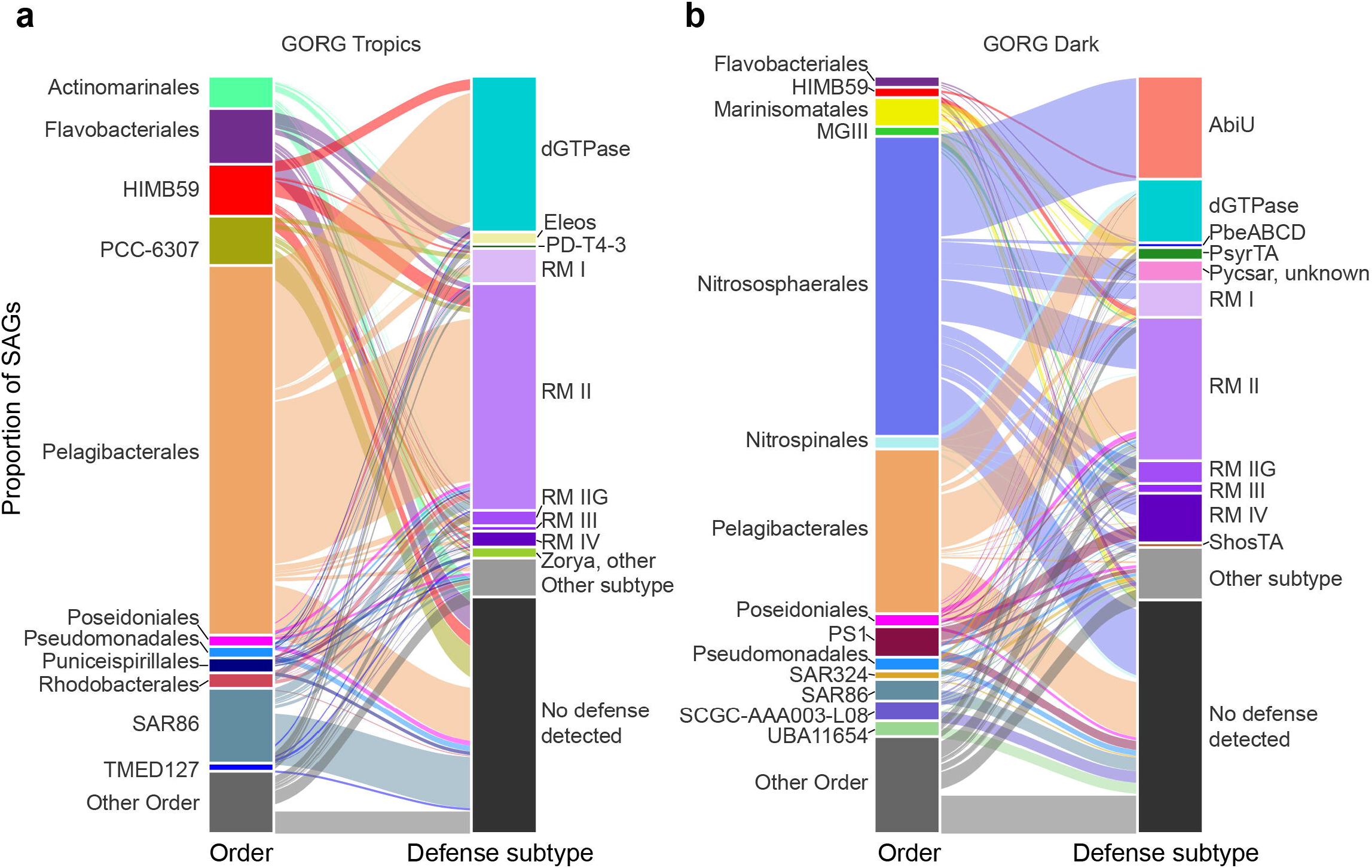
Composition of taxonomic order and defense subtypes in GORG Tropics (a) and Dark (b). Left strata display the proportion of SAGs in an order with same hued lines connecting SAGs to the defense subtypes they encode in the right strata. The right strata are the proportion of SAGs with that defense subtype. “Other order” refers to SAGs belonging to orders that corresponded to < 1% of the prescribed dataset. “Other subtype” refers to subtypes that were present in < 1% of SAGs of the prescribed dataset. “None” subtype refers to SAGs with no defenses.

Further illustrating the prevalence of RM II, 75% of SAGs that contained more than one defense (31% of GORG in total) encoded at least one RM II system. Overall, the co-occurrence of system subtypes reflected the general abundances of systems (Supplemental Fig. S3). This lack of distinctive system co-occurrence contrasts surveys of reference genomes, in which combinations of defense system types or target specificities have been shown to consistently co-occur and enhance or complement each other, resulting in a broader host resistance ^29,30^.

The absence of detectable system relationships here may stem from the fact that most cells only encoded one defense.

Given that a diverse majority of GORG SAGs were defended by RM II (often as their only predicted defense), we speculated that RM II genes would exhibit high sequence variation to widen the viruses a host population can target, suggested by the pan-immunity model ^10^. To assess this, we separately compared the amino acid sequences of the two proteins necessary for RM II function: RM II restriction endonuclease (REase) and RM II methyltransferase (MTase) genes. Alphaproteobacteria lineages (orders: Pelagibacterales, HIMB59, Rhodobacterales, Puniceispirillales) dominated the RM II-containing genomes (75%) and exhibited significantly higher levels of sequence similarity and phylogenetic congruence for the genes across this class than non-alphaproteobacteria lineages (Fig. 3). For instance, RM II genes from alphaproteobacterial within the same species exhibited significantly higher amino acid identities (average: MTase 98%, REase: 98%) than within non-Alphaproteobacteria species (average: 54% REase; Mtase 49%), and similar patterns were observed at higher taxonomic levels (within a genus, within a family; Supplemental Fig. S4; Supplemental Table S12; Mann Whitney U test, two-sided, p value < 0.05). Furthermore, in phylogenies of REase and MTase, we observed two contrasting patterns. The most abundant species in GORG Tropics was an alphaproteobacterial group *Pelagibacter sp0032099* (2.2% of GORG Tropics), and 70-80% of its members clustered monophyletically in both gene trees. The remaining were scattered throughout, indicating putative lateral gene transfer. Meanwhile, the most abundant species in GORG Dark, the archaeon *Nitrosopelagicus sp000484935* (4% of GORG Dark), clustered with bacterial lineages in both gene trees. The largest monophyletic clade of this species only contained ∼10% of its members’ RM II genes. These observed inter-domain clusters broaden the pan-immunity model beyond closely related species to the sharing of defenses across domains^10^. Notwithstanding, the limited diversity of RM II genes in alphaproteobacterial lineages suggest differential phylogenetic constraints on contributions to a community’s defense pool that potentially reflect unknown, multifunctional roles of RM systems, such as addiction modules ^31^.

**Fig. 3.**
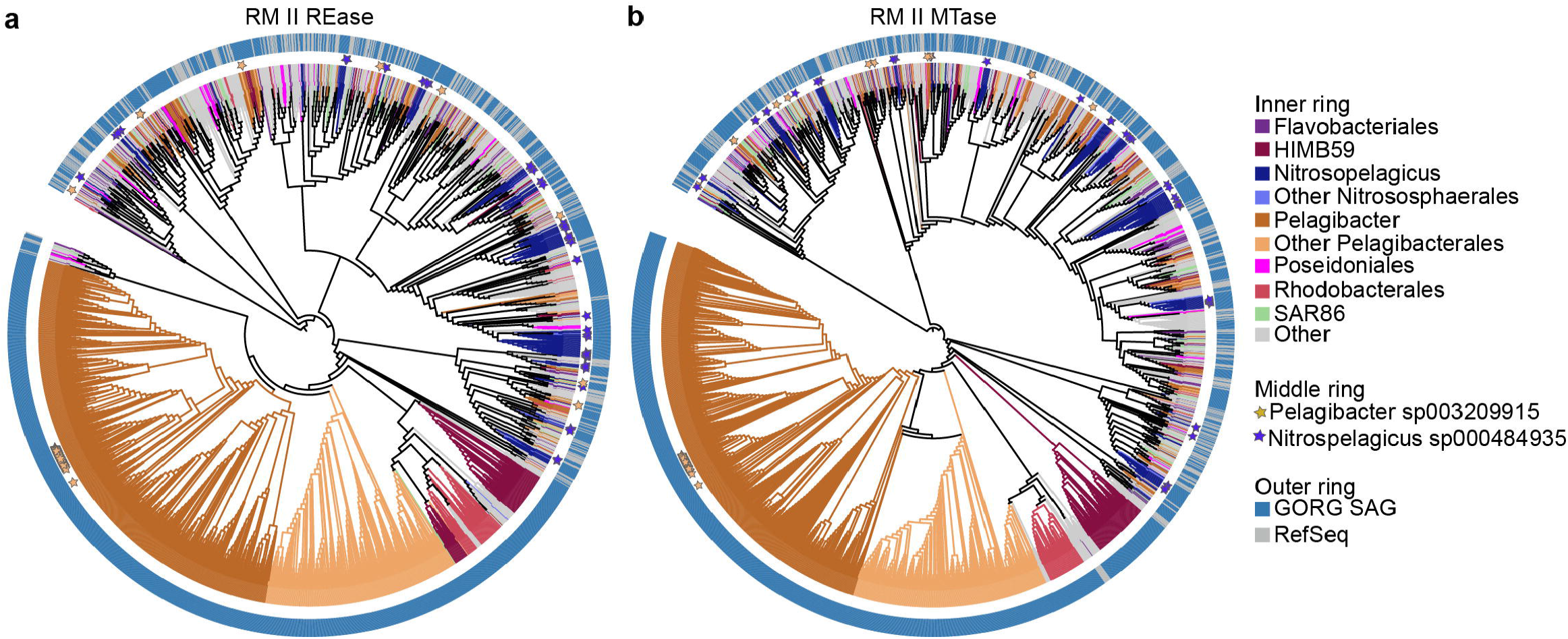
Diversity of defense system subtypes and RM II genes. Phylogeny of RM II REase (a) and RM II MTase (b) amino acid sequences found in GORG and RefSeq study (Tesson et al 2022). Branches and inner ring is colored by taxonomy. Middle ring contains stars indicating the most abundant species in GORG Tropics (orange) and GORG Dark (purple). Outer ring indicates reference versus RefSeq.

### Defenses in abundant lineages of the sunlit and dark oceans

The next most abundant system in GORG Tropics and Dark differed from each other, with the deoxyguanosine triphosphatase (dGTPase) system in GORG Tropics (31% of SAGs) and the abortive infection system AbiU in GORG Dark (21% of SAGs) (Fig 4). The dGTPase system is an enzyme that cleaves dGTP into deoxyguanosine and tripolyphosphate. This enzyme was initially discovered in *E. coli* and likely functions in genomic maintenance ^32,33^. It has recently been found to defend against a range of phages by depleting dGTP in the cell, preventing phage genome replication ^34^. This defense was the second most common defense in our survey of sunlit ocean MAGs (13%), but it is not especially widespread in the broader prokaryotic community (6.7% of RefSeq ^35^). The dominance of dGTPase in GORG Tropics is largely driven by its presence in the order Pelagibacterales (46% of GORG Tropics SAGs, 80% of SAGs with dGTP) (Fig 2a). Most SAGs in this order encoded at least one dGTPase system, in 52% of SAGs across both GORG Tropics and Dark. The dGTPase system has not been previously described in Pelagibacterales genomes, but we also detected this system in reference assemblies of this order in GenBank (Supplemental Table S5, S13). The presence of dGTPase in Pelagibacterales may reflect the genome streamlining strategy of this lineage, as its general function of dGTP cleavage can serve other purposes in the cell ^36–39^. We hypothesize the dual functionality is likely given that this gene was not found near other defense systems (Supplemental Table S14) and instead neighbored genes involved in nucleic acid processing and membrane transport (Supplemental Table S15). Experimental challenges of pelagiphages on *Pelagibacter* cultures would elucidate roles of this gene.

**Fig. 4.**
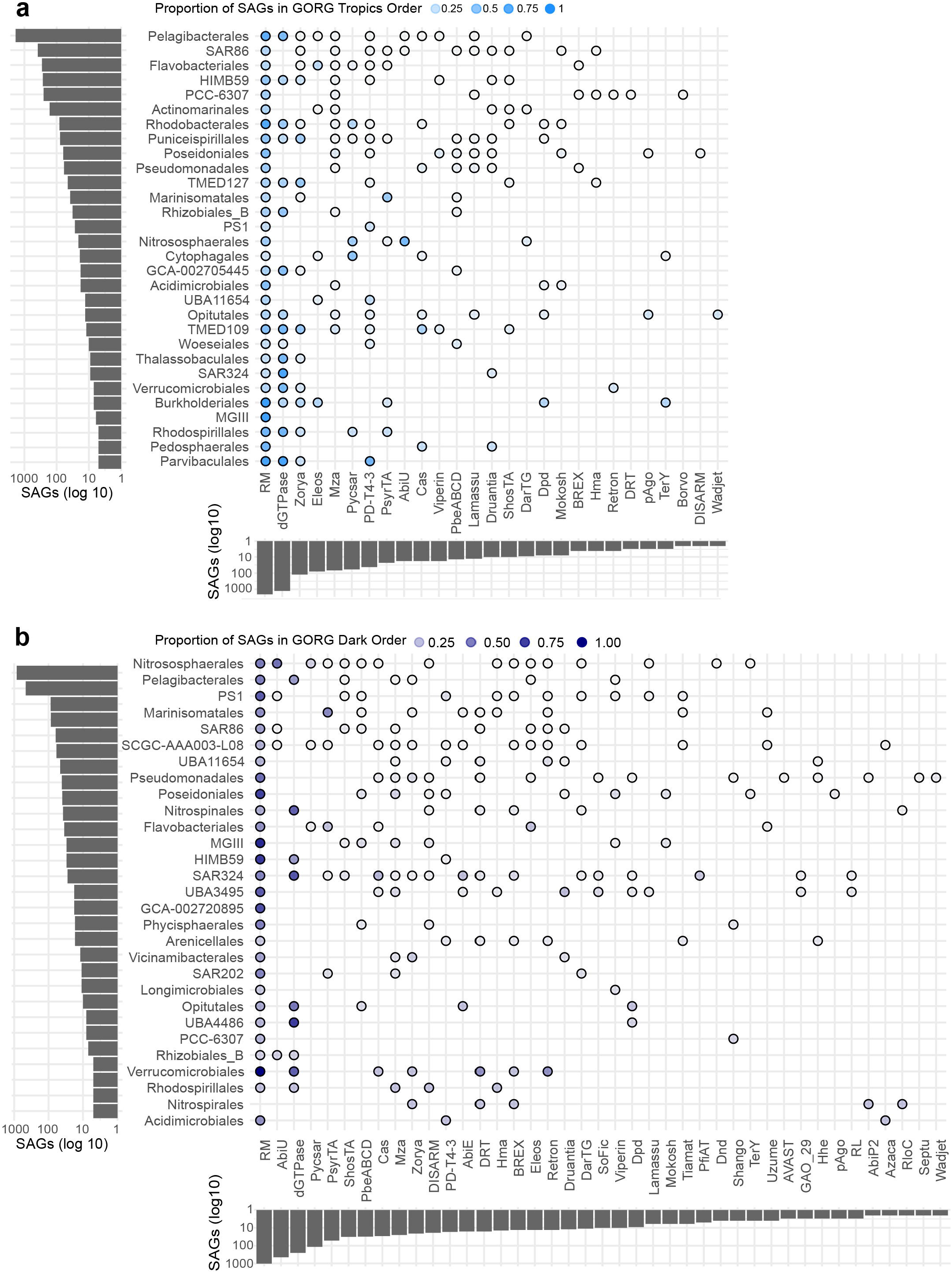
Distribution of defense system types across taxonomic orders in the GORG Tropics (a) and Dark (b). Orders and systems are shown that were present in more than one SAG. The shading of each bubble corresponds to the proportion of SAGs in a given order with that defense system. Y-axes are ordered by SAG abundance and x-axes are ordered by system type abundance.

In GORG Dark, the second most common defense was the AbiU system (Fig 4b) present in 21% of cells (449 SAGs). In contrast, we detected AbiU in only 2% of dark ocean MAGs, and this system is in only 4% and 3% of RefSeq Bacterial and Archaeal genomes, respectively ^35^. Roughly 97% of the SAGs that encoded AbiU belonged to the archaeal order Nitrosophaerales, with the remaining six SAGs spanning five orders in three phyla (Proteobacteria (4), Marinisomatota (1), Chloroflexota(1)). Enrichment of AbiU in abundant dark ocean archaea may stem not only from the dominance of Nitrosophaerales (40% of GORG Dark), but also from the environmental conditions of the dark ocean. AbiU is a type of abortive infection (Abi) system that was initially characterized in *Lactococcus lactis*^40^. Though the details of this mechanism are unknown, experimental work has shown this system sense phage transcription and then stalls cell metabolism, preventing the reproduction and spread of the phage in to other members in its population ^40^ ^41^.

Abi defenses are considered altruistic due to the cost incurred by the infected cell in an effort to protect others. The maintenance of altruistic defense strategies is predicted to be favored in environments where closely related kin are nearby, such as a highly structured environment like biofilms or the human gut, preventing spread of the virus to its relatives nearby. This does not reflect general ecological conditions of the dark ocean, where cell densities are low and populations are relatively well-mixed ^22^. However, modelling work has shown that programmed cell death, like Abi, can be maintained in a well-mixed population over evolutionary time in cases where the cell has a negligible chance of dividing again relative to the viral infection load ^42^, insinuating a slow division rate. The average doubling time for Nitrososphaerales estimated *in silico* here was 12.6 hours, which was significantly higher than the other lineages in GORG Dark (average 11.9 hours; Wilcox test p value < 0.05), but no deep ocean Nitrososphaerales members are in culture to validate this estimate. Another factor favoring altruistic defenses is low viral pressure^11,42^, which does reflect conditions of dark ocean Nitrososphaerales. Viral and cellular concentrations are lower in the dark ocean than the surface ocean ^22,43^; additionally, we detected a lower infection rate of Nitrososphaerales cells (0.7% of SAGs) compared to the GORG Dark average (3.9% of SAGs). Finally, the next common defense in Nitrososphaerales beyond RM (49.9% SAGs) was another altruistic system, Pycsar in 14.7% of SAGs in this order. While culture-based experiments are necessary to confirm the realized functions of the AbiU and Pycsar systems detected in this order here, these results suggest this lineage relies heavily on programmed cell death to combat viruses.

Taken together, the specific enrichment of dGTPase and AbiU in abundant GORG lineages of each depth, (Pelagibacterales and Nitrososphearales, respectively), may reflect the disparate ecological and evolutionary strategies of these lineages, with genome streamlining^36^ resulting in the reliance of Pelagibacterales cells on a multi-functional enzyme for defense (dGTPase), while dark ocean Nitrososphaerales cells rely on an altruistic strategies (AbiU and Pycsar) potentially due to its limited pathogen exposure and slow growth rates^42^.

### Relationships between system diversity and prevalence

The remaining defense systems were relatively rare, detected in a mean of 120 SAGs (1.8% of GORG) and a median of 5 SAGs (< 0.1% of GORG), with 21 of these systems (36%) found in a single SAG. This rarity held true when considering GORG Tropics (median 0.16% (7.5) SAGs) and GORG Dark (median 0.19% (4) SAGs), separately. Despite the general rarity of systems in the ocean, non-singleton defense systems were broadly distributed across distant taxa. Roughly 90% of these systems were found in at least two phyla. Overall, system types were found in a median of 4 phyla, 4 classes, 7 orders, 8 families, 9.5 genera, and 7 species (28% of SAGs could not be classified to species-level; Fig 5a), suggesting a lack of overall taxonomic specificity. For a given system, the number of SAGs with that system was positively correlated with the number of taxa with that system. This correlation held true for all taxonomic ranks, and was statistically significant (Fig 5b; Kendall correlation *p values* < 0.01; Supplemental Table S16), which indicates a system’s taxonomic distribution is tied to its general prevalence and not tied to the abundance of a particular lineage. Likewise, the number of SAGs in a taxon significantly positively correlated with the diversity of systems in the taxon at all levels (Fig 5c; Kendall correlation p values < 0.01; Supplemental Table S17 for non-order results). This trend suggests that lineages are indiscriminately acquiring defense systems. We suspect they acquire them laterally rather than vertically given their lack of conservation and retention, as most systems were only found in a few SAGs per lineage (Fig 4). Collectively, these results suggest that defenses can be found in a range of lineages rather than exhibiting taxonomic specificity. Moreover, lineages are flexible in which defenses they acquire, but few of these systems are ultimately retained and conserved given their general rarity.

**Fig. 5.**
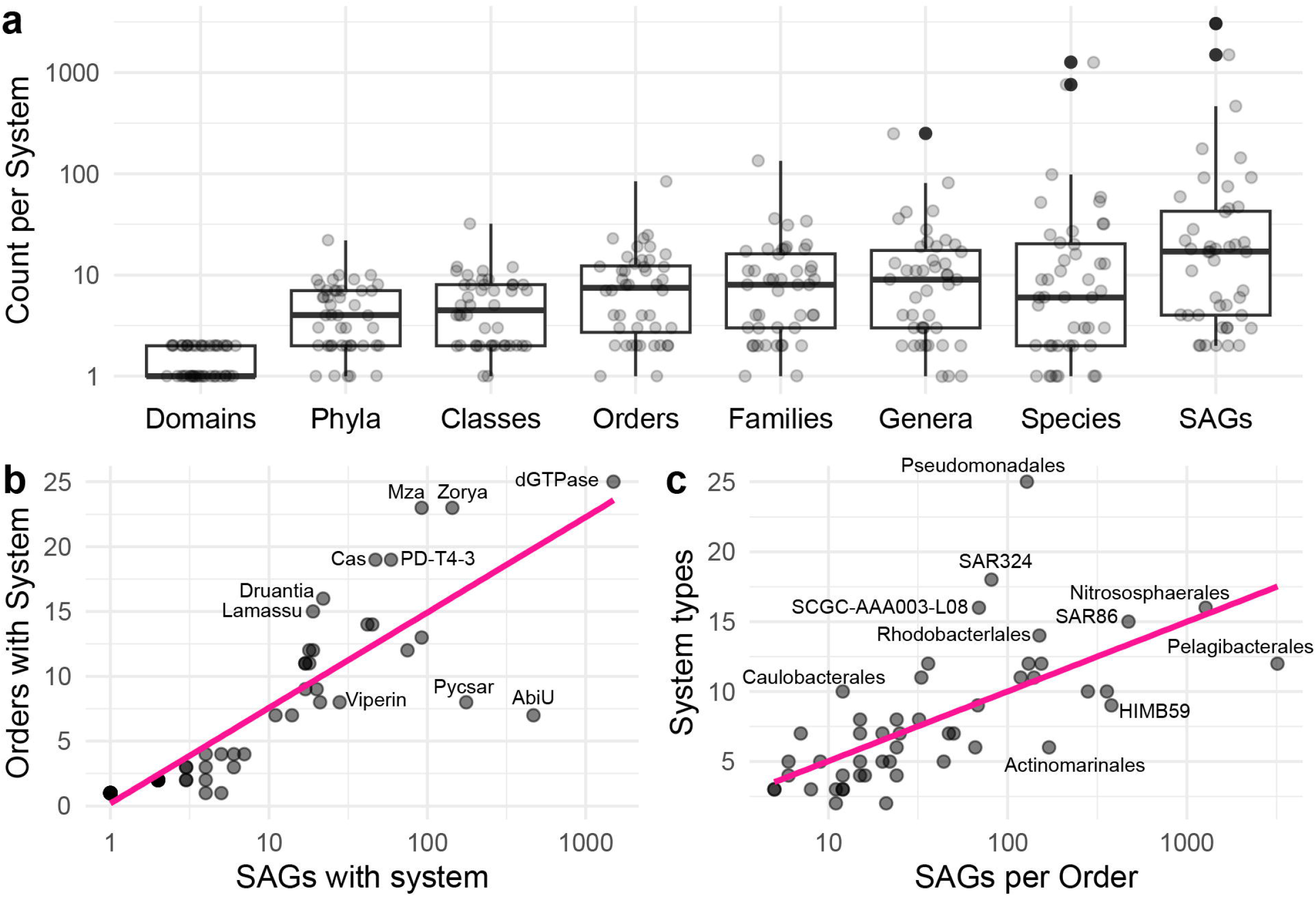
**Relationships between the system or taxonomic diversity versus their abundance**. (a) The number of lineages at a given taxonomic level for each system with at least two SAGs. Median in black midline. Each system is a transparent point. (b) Correlation between the number of orders with a given system versus the number of SAGs. (c) Correlation between the number of types of systems found in an order versus the number of SAGs in that order. (a,b) X-axes are log-transformed. Pink line indicates linear regression. RM is omitted for visibility. System or taxonomic orders are labeled when possible. Shaded area corresponds to linear regression standard error.

Due to the biotechnological implications of CRISPR Cas (Cas) systems, we further examined Cas system occurrence in GORG with a more thorough detection pipeline using CRISPRCasFinder^44^ and CRISPRCasTyper^45^ (see Methods; Supplemental Table S18). This yielded a much lower detection of Cas systems, with only ten Cas systems found across ten SAGs of seven taxonomic orders, versus 56 SAGs across 19 orders in our initial detection. This limited detection is in line with previous studies showing Cas systems are less common in environments with high viral diversity like the ocean ^46^, as well as less common in abundant marine genera ^47^, which SAG recovery tends to favor abundant lineages ^16^. The paucity of Cas systems here also highlights the difference in composition of defenses between references and environmental genomes, as Cas is the second most common defense in prokaryotes on RefSeq, found in 38% of genomes ^35^.

## Conclusion

We present the composition of immune systems in prokaryoplankton from the surface to the deep ocean at single-cell resolution with over 6,000 SAGs from the GORG collection^20^. We found most cells contain at least one defense, with similar overall abundance in defense systems of prokaryotes in the sunlit surface and the dark ocean interior, despite vast environmental and taxonomic differences. We found that prokaryoplankton lineages tightly control their genomic real estate for defenses, regardless of their abundance, estimated growth rates, and estimated viral infection rates. These results suggest that phylogeny must be considered when calculating the resistance parameter in modeling virus-host dynamics ^6^.

Furthermore, incorporating lateral gene transfer rates may be essential, as we found evidence for the cross-domain exchange of the most common defense system, RM II, between archaea and bacteria. Additionally, many rare systems were found sparsely across distant taxa, although cases of high congruency between in defense gene phylogenies with taxonomy were also identified in some lineages. These results suggest the pool of defenses available to a community varies among lineages and, in some cases, extend beyond closely related individuals, as initially proposed in the pan-immunity model ^10^. For instance, we observed RM II systems of alphaproteobacterial groups had higher genetic similarity and phylogenetic congruence than within non-alphaproteobacterial lineages, indicating a smaller pool of defenses for alphaproteobacteria relative to non-alphaproteobacteria groups. Finally, the obtained evidence of phylogenetically broad, including cross-domain, exchange of RM II genes improves our understanding of the biology of this prevalent antiphage defense and also calls for caution when using methylation patterns to link phages with their hosts, as applied in recent epigenomic studies ^48,49^.

Although we found that a genome’s defense abundance is not readily explained by potential for viral contact (abundance, growth rate, estimated viral infection rate, depth), defense-type distributions hint at hosts’ ecological strategies. For example, a multifunctional enzyme (dGTPase) is prevalent in the most abundant sunlit ocean order Pelagibcterales, which may reflect genomic streamlining in surface ocean microbes^50^, given the versatility of this gene ^39^. Whereas in the dark ocean, cells of the most dominant order Nitrosopumilales encode altruistic defenses (AbiU, Pycsar) ^40^ ^11,22^, that may reflect a sufficiently low viral pressure relative to its division rate that renders this strategy more useful than non-fatal immune systems ^42^. Future work targeting defenses in marine prokaryotes, such as cloning candidate defense genes into model systems from metagenomic selections^51^, could assist in refining the role of ecological factors in defense system strategies. This approach could also reveal novel defenses currently undetectable by testing other genes with unknown functions that may shed light on defense distribution trends not possible to observe presently. Collectively, this study provides a foundation for incorporating antiphage resistance of prokaryoplankton into estimating impacts of the viral shunt on biogeochemical cycles across the ocean.

## Acknowledgements

We thank the GORG Consortium for early access to the GORG-Dark dataset and for advice. We thank Luis M. Bolaños for discussion on Pelagibacterales diversity.

## Code availability

All Nextflow, R, and Python scripts for calculating statistical tests and generating figures (except for the phylogenies, see Data availability) are available on the GitHub repository GORGdefenses (https://github.com/scubalaina/GORGdefense).

## Data availability

The GORG Tropics SAG data was downloaded from NCBI BioProject: PRJEB33281 and the GORG Dark SAG reads are available on NCBI SRA BioProject: PRJNA1259050, and the assemblies used on FigShare: https://figshare.com/s/59f42e6ffe5629d6fd79. The data and files for generating and annotating the phylogenetic trees are on Zenodo (DOI: 10.5281/zenodo.22086263). All Supplemental Tables (S1-S18) are sheets in the excel file Supplemental Dataset 1.

## Methods

### SAG curation and defense detection

Nucleotide sequences and sample information of single-amplified genomes (SAGs) were compiled from the Global Ocean Reference Genome collection (GORG) Tropics ^20^ and GORG Dark^21^. SAGs were filtered for those with at least 50% completeness. The GORG Dark SAGs sourced from the Arctic, Ross Ice, Black Sea, Baltic Sea niches were excluded, as these communities were determined as disparate from the pelagic open ocean community^21^. Amino acid sequences were predicted and GFF files were prepared of the GORG SAGs with Prodigal (v 2.6.2 ^52^). Defense systems and genes were detected using defense system detection tools DefenseFinder (v1; ^9^) and PADLOC (v 1.1.0; ^23^). DefenseFinder was run with default settings on the SAG proteins. We ran PADLOC on SAG proteins and gff files, using default parameters aside from the flags --fix-prodigal --outdir. Since some defense systems were unique to each tool, we used both outputs in total. There were some discrepancies in the tools in terms of labeling and capitalization (e.g. “RM_Type_II” in DefenseFinder is “RM_type_II” in PADLOC). Additionally, PADLOC only indicated the subtype of a system, and we thus adopted the system-level designation used by DefenseFinder when available, and we relabeled the systems to use consistent formatting (Supplemental Table S3). Moreover, PADLOC contained a defense system called “DMS” for cases where any two proteins of a DNA modification system (DMS) such as RM or phosphothioration were detected, but both did not correspond to a canonical system (e.g. an RM endonuclease with a PdbE gene). To be conservative in determining the frequency of a single type of system in a SAG, we only counted a unique instance if none of the proteins overlapped between different instances in the tool. For example, in SAG AG-538-P13, PADLOC detected two RM systems [RM_2: AG-538-P13_NODE_2_38,39,40,42 | RM_3: AG-538-P13_NODE_2_42] and DefenseFinder detected two RM systems [RM_1: AG-538-P13_NODE_2_39,40 | RM_2: AG-538-P13_NODE_2_42]. These would all be counted as one instance of an RM system since at least one protein was found to overlap with all sets. This leads to a potential underestimate of system frequency, but the lack of experimental validation for these genes overall warrants conservative estimates of occurrence. To perform this parsing and consolidation, we used a custom python script (parse_merge_dfploc.py). Detected defenses systems and subtypes are reported in Supplemental Table S1 and S2, respectively.

### MAG and RefSeq comparison

Metagenome-assembled genomes were obtained from the study by Paoli et al. 2022 ^53^. We filtered these MAGs to retain those with at least 50% completion and a maximum of 10% contamination according to CheckM (v1) ^54^. Those corresponding to the sunlit ocean were selected by Chang et al. 2024 ^16^ (Supplemental Table S5). Those corresponding to the dark ocean were selected from the remaining MAGs belonging to samples collected below 200 meters according to Paoli et al 2022 (Supplemental Table S4). In total, we retained 10,056 MAGs. Defenses were detected with PADLOC and DefenseFinder as described for the SAGs (Supplemental Table S5). For the RefSeq defenses, genome assembly information and defense detection results were downloaded from the RefSeq DB page of Defense Finder Webservice ^35^.

### RM II gene diversity

Amino acid variation of RM type II restriction endonuclease (RM II REase), and RM II methyltransferase (RM II MTase) was assessed via BLASTp. First, SAG proteins encoded by these genes were aligned against each other using BLASTp (E value < 10^−6^). The amino acid percent identity was retained between each pair of hits based on their best alignment by highest bit score, since multiple alignments between the same proteins can occur. To reconstruct phylogenies of these genes within a broader prokaryotic diversity, we compiled the accessions of proteins from RefSeq DB on the Defense Finder Webservice ^35^ and downloaded their sequences via Batch Entrez on the NCBI. To refine this reference dataset to a tractable number of proteins, we retained only reference proteins that were best matches to GORG proteins via BLASTp (E value < 10^−6^) based on the highest bit score. Likewise, we filtered GORG sequences to remove identical sequences, via cd-hit (-c 1; v4.7 ^55^). After these filtering steps, remaining proteins were aligned with Muscle5 ^56^. Alignments were trimmed for those with more than 90% gaps using trim-Al (-gt 0.1) ^57^. Phylogenies were reconstructed with IQTREE (-m TEST -wbt -bb 1000 --runs 5 -safe; version 2.2.5; ^58^). Trees were visualized in iTOL (v7 ^59^) and layered with iTOL-formatted annotation files. All amino acid sequence files, BLAST results, alignments, and treefiles are available on the Zenodo repository.

### Virus infection detection

All contigs over 10 kilobases within a SAG were run through VirSorter2 ^60^ as 10 kilobases results in the highest accuracy of this tool ^61^. Sequences were retained as viral if they had at least two viral hallmark genes detected by VirSorter2 ^62^. The presence of a putative viral contig in a SAG was interpreted as an infection (Supplemental Table S1, Table S11).

### Visualizations and statistics

All figures aside from the phylogenies were generated with R in Rstudio and refined in Adobe Illustrator (2026, v. 30.7), using the packages ggplot2, ggpubr, and ggalluvial. Statistical tests were run in python using the modules scipy, numpy, and pandas (Python 3.11). P values were reported in supplemental tables and significance was considered for values below 0.05. All code and related files are available on the GitHub repository GORGdefense.

### CRISPR Cas detection and analysis

We searched for CRISPR Cas systems in the GORG data using CRISPRCasTyper ^45^, with default parameters in meta mode (--prodigal meta) and keeping temporary files (--keep_tmp). The results included systems which are confidently predicted (‘True’), low quality (‘Putative’) and Cas proteins with missing CRISPR array (‘Orphan’). Contigs with CRISPR Cas systems were extracted and entered into the CRISPRCasFinder ^44^ web service (with default parameters) in order to detect CRISPR arrays, direct repeat (DR) and spacer sequences. If ‘Orphan’ systems were predicted by CRISPRCasFinder to contain a CRISPR array, then we retained them for further analysis. We used CRISPRCasTyper to create a database of CRISPR Cas systems found in GORG data. This database recorded (1) the SAG and contig in which each putative CRISPR Cas system was found, including its predicted class, type, and subtype (where available), as well (2) as the Cas operon start and end positions in the contig, and (3) the DR and spacer sequences predicted by CRISPRCasFinder (Supplemental Table S18).

## Abbreviations

GORG: Global Ocean Reference Genome
SAG: single-amplified genome
MAG: metagenome-assembled genome
RM: restriction modification

