## Supplemental Figures for "Global, single-cell-resolution of antiviral defenses in marine prokaryoplankton"

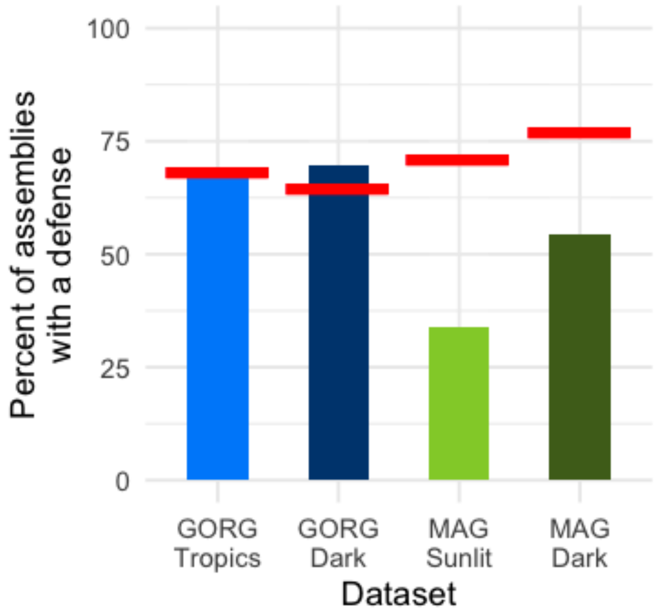


**Supplemental Figure S1**. Percentage of genome assemblies with a defense across dataset types (GORG SAGs versus MAGs) versus their average genome completion (red bars). Colors distinguish datasets.

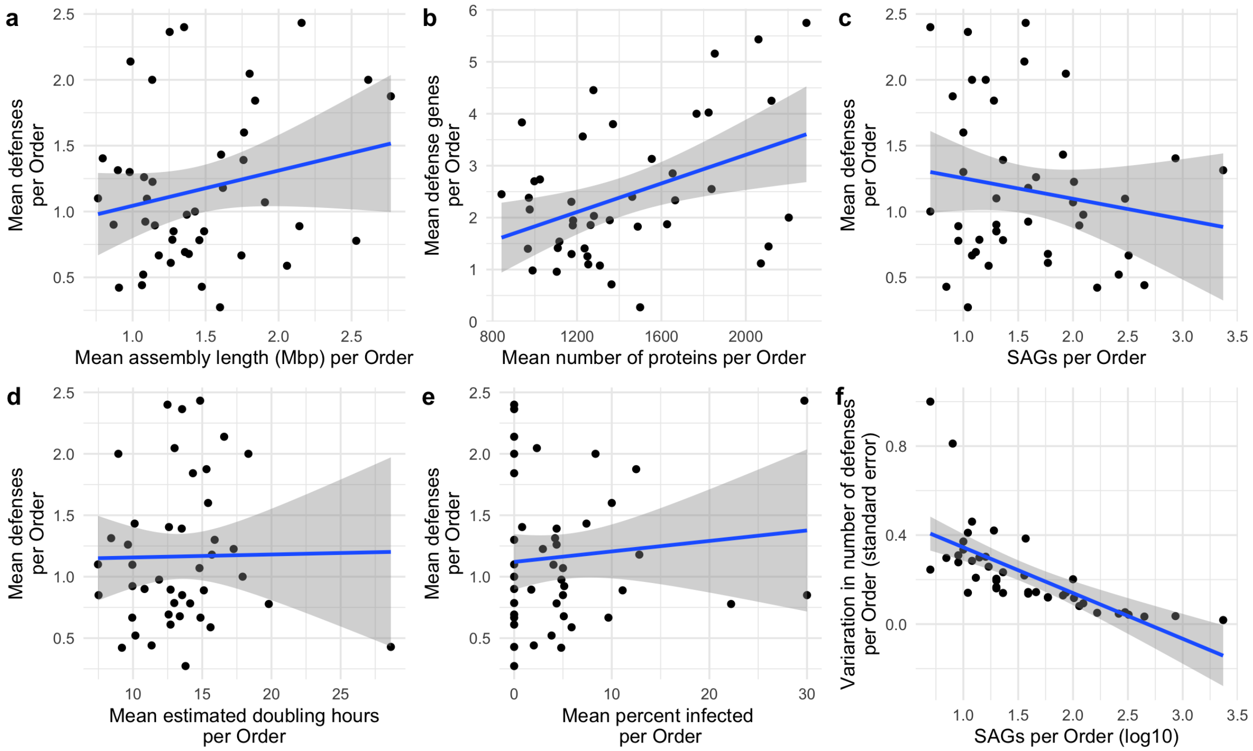


**Supplemental Figure S2.** Relationships between number of defenses (a,c,d,e) or defense genes (b) versus ecological and genomic factors per Order in GORG. (f) Variation in the number of defenses per Order via standard error versus Order abundance via number of SAGs.

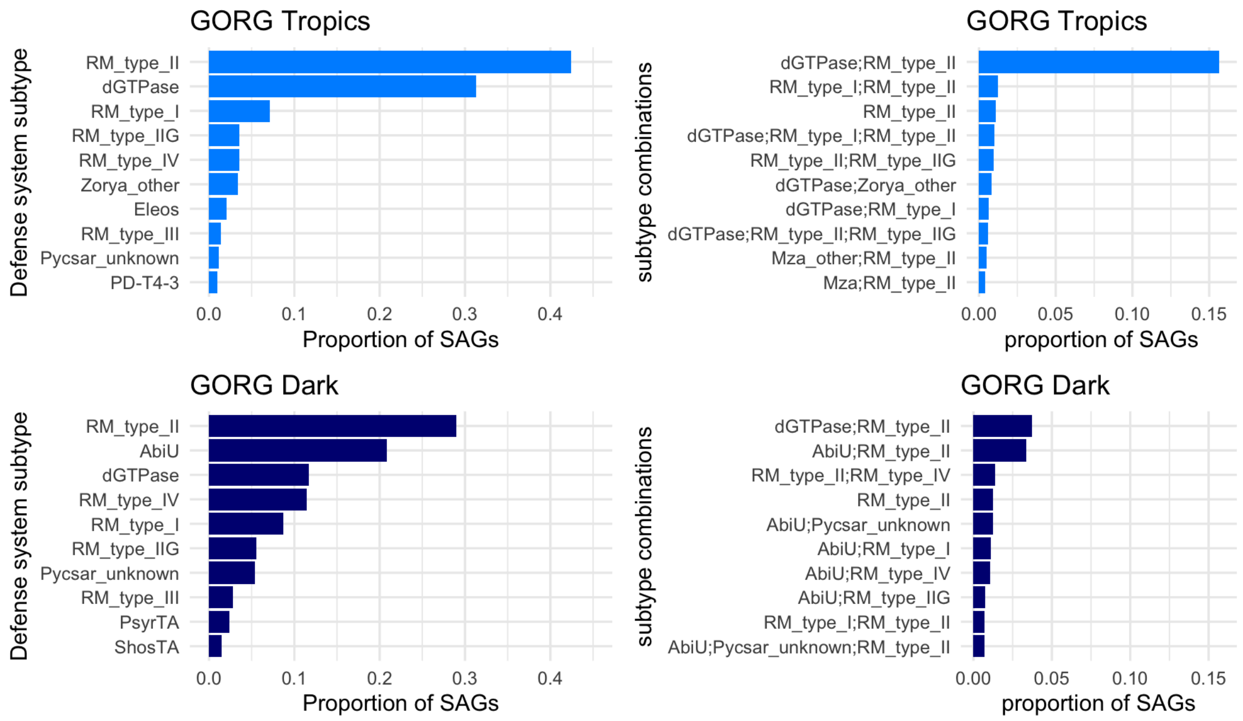


**Supplemental Figure S3.** Proportion of SAGs with a given defense subtype or combination of defense subtypes.

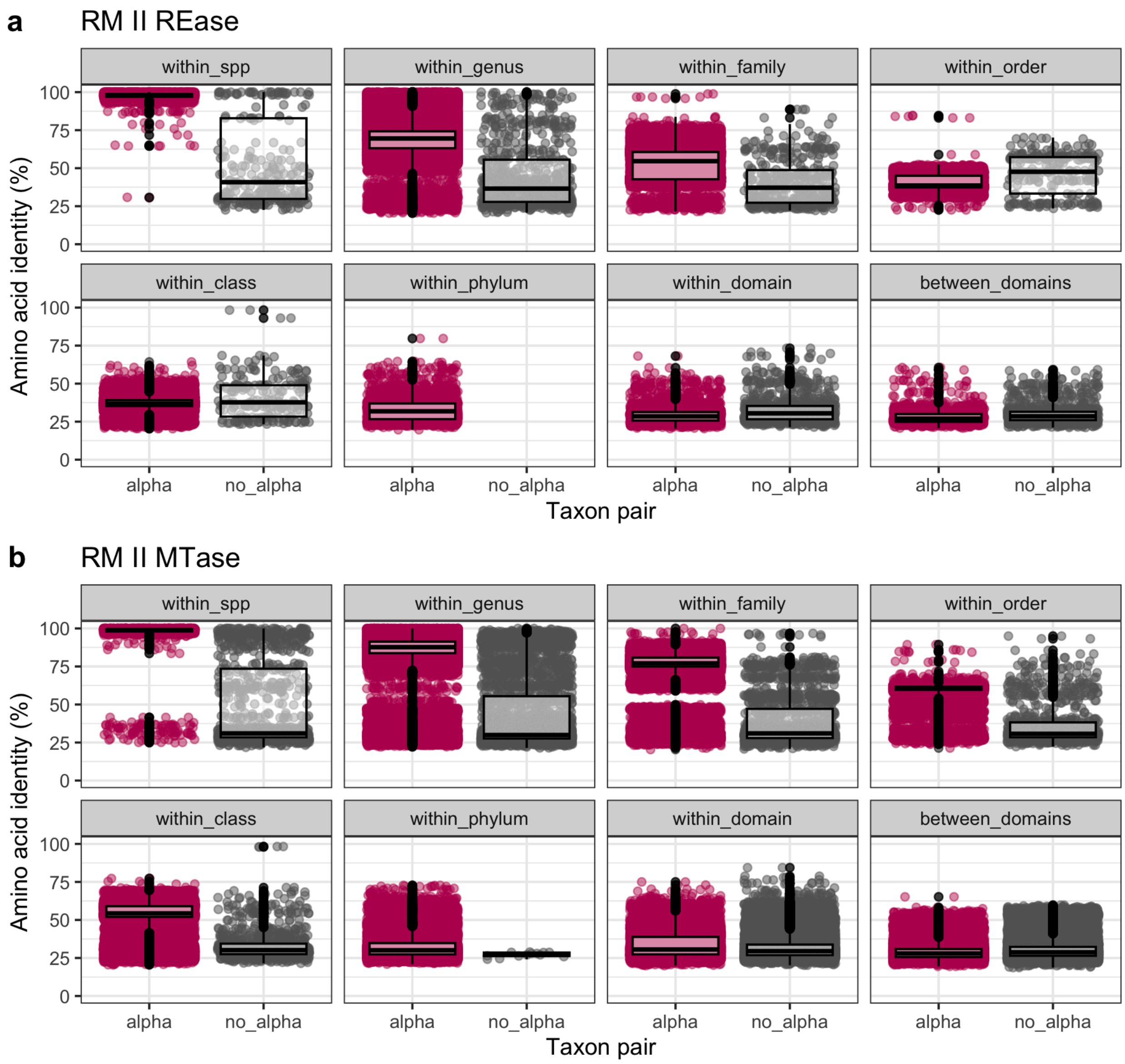


**Supplemental Figure S4.** Amino acid identity of RM II REase (a) or MTase (b) genes of different taxonomic distances (e.g. within species, within genus but different species, etc.) Lineages that contain alphaproteobacteria are distinguished from other lineages on the x-axes and with maroon jitter points.
